# Tyrosinase mediated humic substances synthesis by *Bacillus aryabhattai* TFG5

**DOI:** 10.1101/322024

**Authors:** Iniyakumar Muniraj, Syed Shameer, Priyadarshini Ramachandran, Sivakumar Uhandi

**Affiliations:** Biocatalysts lab, Department of Agricultural Microbiology, Tamil Nadu Agricultural University, Coimbatore, Tamil Nadu, India

**Keywords:** *Bacillus aryabhattai* TFG5, Tyrosinase, Laccase, coirpith biomass, oxidative polymerization, humification

## Abstract

The present investigation aims at understanding the mechanism of Humic Substances (HS) formation and enhancement through tyrosinase produced by *Bacillus aryabhattai* TFG5. A bacterium isolated from termite mound produced tyrosinase (1.34 U.ml^−1^) and laccase (2.1 U.ml^−1^) at 48 and 60 h of fermentation respectively. The protein from *B. aryabhattai* TFG5 was designated as TyrB and it had a predicted molecular weight of 35.23 kDa. Swiss modelling of protein revealed a bi copper protein with its conserved residues required for activity. Interestingly, TyrB efficiently transformed and polymerized standard phenols besides transforming free phenols of Coir pith Wash Water (CWW). In addition, spectroscopic evidences suggest that TyrB enhanced the HS production from coir pith biomass. Furthermore, degradative products and changes in biomass structure by TyrB analysed through FT-IR suggests that TyrB might follow the polyphenol theory of HS synthesis.

## Introduction

Tyrosinases(monophenol, *o*-diphenol:oxygen oxidoreductase,EC 1.14.18.1) are bifunctional type 3 copper containing enzymes which catalyze-hydroxylation of monophenols yielding diphenols(cresolase activity) and subsequent oxidation of *o*-diphenols to quinones (catecholase activity)[1, 2]. Tyrosinases are widespread among plants, animals, fungi, and bacteria[3]. They are essential in melanin biosynthesis and also used in pulp and paper manufacturing [4], textile, pharmaceutical industry[5]. In addition, tyrosinase is also efficient in oxidizing low molecular weight phenols and finds a suitable place in wastewater treatment [6].

Under soil environments, tyrosinase is believed to participate in enhancing the formation rate of humic substances (HS)[7]. Such HS are ubiquitously obtained by oxidative biotransformation of dead organic matter in the soil. The resident time of HS in soil is 10^2-^10^3^ years and thus the humic substances formation (humification) is considered as one of the key processes in atmospheric carbon dioxide sink. It is estimated that 1462 Pg of Carbon is found in the total biosphere[8], of which about one third (470 Pg) of carbon is found the soil. Therefore increasing the rate of humification is important for long time storage of carbon in soil [9]. Besides, humification indirectly reduces global warming by sequestering the atmospheric CO_2_ in soil [10].

HS, being largest pool of recalcitrant organic carbon in the terrestrial environment, its formation, oxidative biotransformation and mineralization in soil is predominantly due to the oxidative enzymes mostly from the fungal origin. Lignin peroxidase (LiP), Mn-dependent peroxidase (MnP), versatileperoxidase (VP), other peroxidases, laccase, and tyrosinase are the major oxidative enzymes involved in the formation of HS[9, 11]. Among them, tyrosinase catalyses the oxidation of a phenolic and non-phenolic portion of the substrate into quinones and aryl radicals respectively. HS formation by fungal tyrosinase in wood and soil has been studied previously[7]. Several tyrosinases from bacteria have been described. For instance, tyrosinase from *Streptomycessp*. has been described widely[12]. Other bacterial sources of tyrosinase such as *Aeromonas media*[13], *Azospirillum* sp, *Bacillus megaterium*[14], *Bacillusthuringiensis*[15],*Marinomonas mediterranea*[16],*Pseudomonas*[17], *Rhizobium meleloti*[18],*Thermomicrobium roseum*[19]and *Verrucomicrobium spinosum*[20]have also been described.

However, the bacterial tyrosinase described above are used for specific applications such as solvent tolerance, water decontaminants, phenol removal, detoxification of plant host defences, thermal and salt tolerance etc. Studies related to bacterial tyrosinase and its involvement inHS formation are very sparse in literature and it is important to elucidate the mechanisms of HS formation by bacterial tyrosinase. Hence, this investigation aimed at understanding the role of tyrosinase from bacterial isolate *Bacillus aryabhattaiTFG5* in HS production. In addition, the enzyme was also evaluated for oxidation of phenols from CWW.

## Materials and Methods

### Culture, materials, chemicals and media

Tyrosinase of *Bacillus aryabhattai*TFG5,a newly isolated bacterium of Termite Fungal Garden by the authors,was used for HS formation experiments[21]. Coir pith biomass and Coir Pith Wash Water were obtained from local industry and used for oxidative transformation studies. 3-methyl-2-benzothiazolinone hydrazine (MBTH), L-Tyrosine, LDOPA, ABTS (2,2’-azino-bis(3-ethylbenzothiazoline-6-sulphonic acid) and mushroom tyrosinase were obtained from SigmaAldrich India (Bengaluru). p-hydroxy benzoic acid, 2, 6, Dimehoxy phenol, p-cresol, p nitro phenol and catechol were obtained from HI-Media Laboratories Pvt. Ltd (Mumbai). Laccase production was monitored in Crawford’s media containing (g.L^−1^) Glucose 1.0, Yeast extract 1.5, Na_2_HPO_4_(4.5), KH_2_PO_4_(1.0), MgSO_4_. 7H_2_O(0.12), NaCl(0.2), CaCl_2_(0.05). Tyrosinase production was evaluated in the medium containing (g.L^−1^) casein broth hydrolysate (10), K_2_HPO_4_ (0.5), MgSO_4_ (0.25), L-tyrosine (1).*B.aryabhattai* TFG5 was maintained in Luria Bertani (LB) broth containing (g.L-^1^) tryptone (10), yeast extract (5), and NaCl (10).

### Time course production of tyrosinase and laccase

One-day-old culture of *B.aryabhattai*TFG5 grown in LB broth having OD value of 0.1 was transferred into respective sterile media (50 ml) in 250 ml Erlenmeyer flasks for monitoring the laccase and tyrosinase production. The flasks were incubated in an incubation shaker (New Brunswick, USA) at 30 °C. Enzyme activity and growth were monitored at 4-h intervals for 68 h by monitoring changes in absorbance at 505nm and optical density at 600 nm respectively in a multimode spectrophotometer Spectra Max 360 (Molecular Devices, USA). Flasks were removed at periodic intervals and the contents were centrifuged in a microfuge (Biorad, USA) and the supernatants were used for enzyme assay.

## Bioinformatics analyses of TyrB

Tyrosinase from *B.aryabhattai* TFG5 was named as TyrB. Gene-specific primers of *B. Megaterium*tyrosinase (Forward 5’-GAGGTTAAACCATGGTAACAAGTATAGAGTTAGAAAAAACG-3’ and Reverse 5’ -TGCTGTTTCTAGATCTGGTTAATGGTGGTGATGGTGATGTGAGGAACGTTTTGAT TTTC-3’) [22] were used to amplify tyrosinase gene of *B. aryabhattai*TFG5 and sequenced at Scigeneome Pvt. Ltd., Cochin, Kerala, India. The gene sequences were translated into protein and were aligned using multiple sequence alignment (MSA) tool in Bio edit version 7.2.5. The protein sequences of various tyrosinase were obtained from NCBI (www.ncbi.nlm.nih.gov) or RCSB (www.rcsb.org) data repositories and the homology model of TyrB was built using automated Swiss-modelling server which was verified using Structure Analysis and Verification Server version 4 (http://services.mbi.ucla.edu/SAVES/)[23]. The homology model of TyrB was visualized using open source PyMOL version 0.97 (2004). The theoretical molecular weight was predicted online using compute Mw tool (http://web.expasy.org).

## Enzyme assay

The laccase activity was determined at 30 °C for 5 min using 1mM ABTS by monitoring change in absorbance at 420 nm (€ max &#x003D; 3.6 × 10^4^ M^−1^cm^−1^) spectrophotometrically in a Spectramax 360 (Molecular devices, USA). The reaction mixture contained appropriately diluted enzyme which was mixed with 1mM ABTS in sodium phosphate buffer (50 mM, pH 4.5)[24]. Tyrosinase activity was determined at 30ºC for 5 minutes using 1.5 mM L-DOPA (ε505 &#x003D; 2.9×10^4^ M^−1^.cm^−1^)[25]. The reaction mixture contained phosphate buffer (50 mM, pH 7), 1.5 mM L-DOPA, 5 mM MBTH (3-methyl-2-benzothiazolinone hydrazone) 2% n-n’-dimethylformamide, 0.1 mM sodium azide and 10 μL of appropriately diluted enzyme. The reaction was stopped by adding 100 μL of 1 M perchloric acid and the absorbance was measured at 505 nm. One unit of enzyme activity was defined as the amount of enzyme required to oxidize 1 μM min^−1^ of the substrate under standard assay conditions.

## Oxidative polymerization of phenolic compounds

Six different phenols including mono (*p*-hydroxy benzoic acid, 2, 6, Dimehoxy phenol, *p-*cresol, *p*-nitro phenol) and di-phenols (Catechol, Levo DOPA) at 2mM were evaluated for oxidative polymerization experiment. Phenols (2mM) were taken into a 250 ml conical flasks containing 50 ml of the reaction mixture in 50 mM sodium phosphate buffer (pH7.0). TyrB (10U.ml^−1^) was added and the flasks were incubated in dark conditions for 48h. Oxidative polymerization of phenols was estimated according to[26, 27]. Furthermore,the contents were centrifuged for 10 min at 2500 rpm in a microfuge (Biorad USA) and the liquid phase was analysed immediately by Fourier Transform Infrared (FT-IR) spectroscopy. The liquid portion (10μl) was applied to the diamond attenuated totally reflexion (ATR) crystal and analysed by JASCO 7000 FTIR. The infrared radiation was employed through the samples to obtain the corresponding spectrum, which was averaged from several data acquisitions. FTIR spectra were acquired in the wavenumber range of 700–4000 cm^−1^ with a resolution of 4 cm^−1^. After each measurement, the crystalline surface was washed with acetone and dried with a soft paper.[26].

## Transformation of Coir Pith Wash Water (CWW)

The transformation was performed using 50 ml of CWW in 250 ml Erlenmeyer flasks under both sterile and non-sterile conditions. Filter sterilization of CWW was performed by passing the CWW through 0.24μ syringe filters (Pall, Bengaluru). Sterilization of CWW was achieved by autoclave (121^0^C 15lbs for 15 min), while non-sterilized CWW was used as a control. TyrB at 10U.ml^−1^in a total volume of 50 mlwas added in all treatments and the flasks were incubated under shaking conditions at 30°C for 48 h. Transformation products were analysed in a Jasco 7000 ATR- FT-IR. Infra-Red spectra were collected in the range from 4000 to 700 cm^−1^ with a resolution of 4 cm^−1^[26, 27]. In addition, maximum absorbance of the CWW was evaluated in multimode microplate reader Spectra max 360 (Molecular devices USA)[6] and the maximum absorbance of 300 nm was used for change in absorbance studies.

## Humification of coir pith biomass

Coirpith biomass was used as a substrate to produce HS using TyrB[28]. Sieved and dried coirpith biomassat a consistency of 5%was incubated in the presence of TyrB 10U.ml^−1^ in 50mM phosphate buffer at pH7.0 for HS formation[8, 29, 30]. At the end of the experiment, the solid and liquid phases were separated by centrifugationfor 10 min at 2500 rpm (Biorad USA). The liquid phase was analyzed immediately and maintained frozen. In contrast, the solid phase was dried at 105°C for 24 h and maintained at room temperature until use. Depolymerization degree and chemical properties of thecompounds in the liquidphase weremeasured by the increment in the absorbance at 450 nm in a spectrophotometer spectra max 360 (Molecular Devices USA) whichis related with the production of HS. TheE270/400, E465/665, E250/365, E280/472, E280/664, and E472/664 coefficients were calculated to identify unique characteristics of the compoundsfound in the liquid phase[28]. The treated and untreated solid samples were analysed using a JASCO 7000 Fourier Transform Infra-Red (FT-IR) spectroscopy with infrared spectra collected in the range from 4000 to 700 cm^−1^ with a resolution of 4 cm^−1^.

## Results and Discussion

### Time course production of tyrosinase and laccase

Biotransformationof HS are largely oxidativeprocesses with wood- and soil-inhabiting microbes being a major driving force due to the extracellular production of oxidative enzymes of the class, phenol oxidases, which include tyrosinase and laccase. In the present investigation, a potential tyrosinase and laccase producing *B.aryabhattai*TFG5 isolated from termite mound was evaluated for TyrB mediated HS formation. Tyrosinase secretion by TFG5started from 20h, gradually increased and reached its maximum activityat48h (1.34 U.ml^−1^) and its growth in tyrosine broth started 4h onwards and reached maximum at 60h. Similarly, laccase activity (0.2 U.ml^−1^) was initiated after 20h of the fermentation following a maximum production reaching at 68h (2.1U.ml^−1^) of the fermentation, while the highest growthwas observed at68h of the fermentation in Crawford’s broth (Fig 1).

**Fig. 1.**
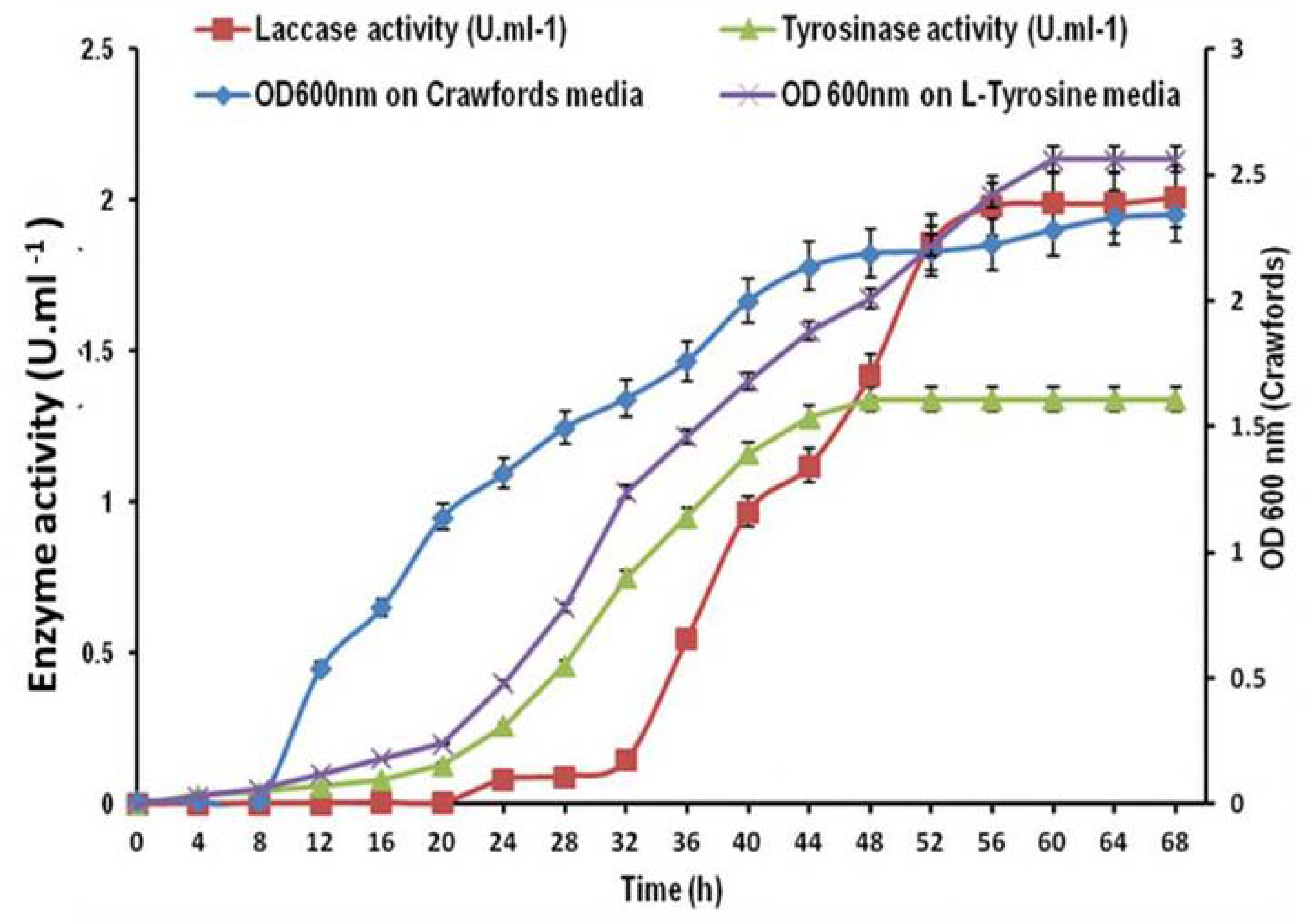
Time course production of laccase and tyrosinase by *B.aryabhattai* TFG5. Enzyme activities were estimated at 4 h intervals on respective media and the optical density at 600 nm measured for the growth of *B.aryabhattai* TFG5. Values are means of three replications and error bar indicates the standard deviation value.

There are few reports on the industrial production of tyrosinase which are generally obtained from fungal sources. Only a few studies to the best of the authors’ knowledge focused on optimization of process parameters for bacterial tyrosinase production. Sambasiva Rao[31]optimized the production conditions of *Streptomyces antibiotics*tyrosinase and showed maximum tyrosinase activity of 4.62 U.ml^−1^. In another study recombinant *E.coli* produced tyrosinase to a high level of 13U.ml^−1^[32]. Comparatively, in the present investigation B. aryabhattai TFG5 produced 1.34 U.ml^−1^ of tyrosinases and when induced for laccase it had produced 1.8 U.ml^−1^without any optimization studies. This suggests that this bacterium would be an advantageous and suitable candidate for HS formation in soil due to its unique property of producing both laccase and tyrosinases.

### Bioinformatics analyses of TyrB

Gene-specific primers of *B. megaterium* were used for amplification of tyrosinase gene in *B. aryabhattai* TFG5 and the ORF encoding the gene was translated into protein. Homology model was predicted inSWISS homologymodelling server using 3NM8 as a template and the structure and thecorresponding metal binding domain are depicted in Fig. 2.

**Fig 2.**
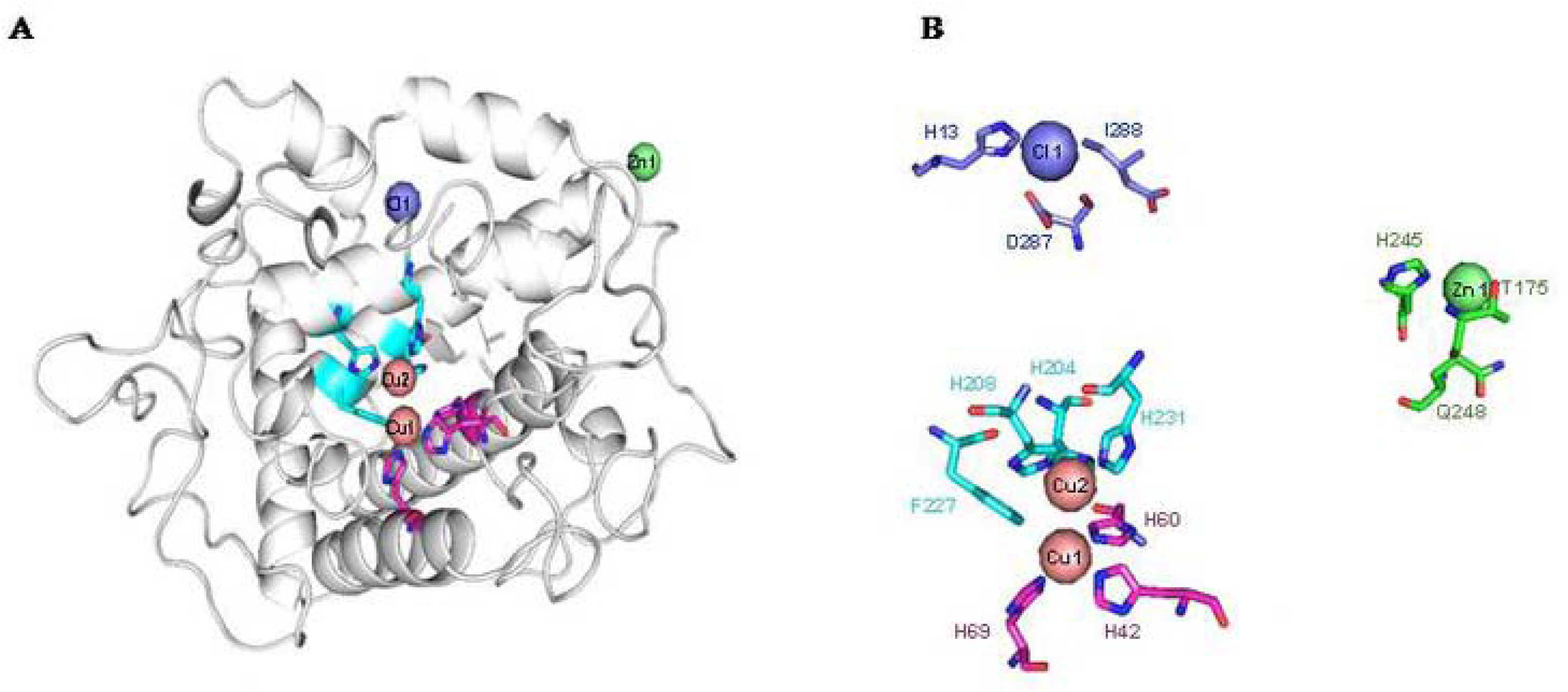
Homology model of TyrB was predicted using crystal structure of *B.megaterium* (3NM8) (A). There were 3 Cu ligands in the model (indicated by sphere) each of which had contact with the His residues and one Phe residue (B).

TyrB had a predicted molecular mass of 35.23kDa and was found to be a bi-copper protein where Cu1 was coordinated with three histidine residues (H69,H42, H60) and Cu2 was coordinated with three histidine (H204,H208,H231) and one phenyl alanine (F227) residues. Other than copper the TyrB also had Cl and Zn binding sites. Multiple alignments of amino acid residues showed that the Cu1 and Cu2 binding residues were conserved with other tyrosinase reported [33](Fig S1). Like most of the bacterial tyrosinases, the TyrB is also a bi copper protein. However, to maintain the structural stability, the enzyme might have coordinated additionally with Zn and Cl ions.

### Oxidative polymerization of phenolic compounds

Monophenols polymerization was relatively higher than diphenols for TyrB as evidenced by the increase in the release of CO_2_ to the former than later. The maximum CO_2_ release was observed in *p*-cresol (477.25 μmol) followed by *p*-hydroxyl benzoic acid (412.5 μmol). The lowest CO_2_ release was observed in Levo DOPA followed by 2,6 DMP (Fig 3).

**Fig 3.**
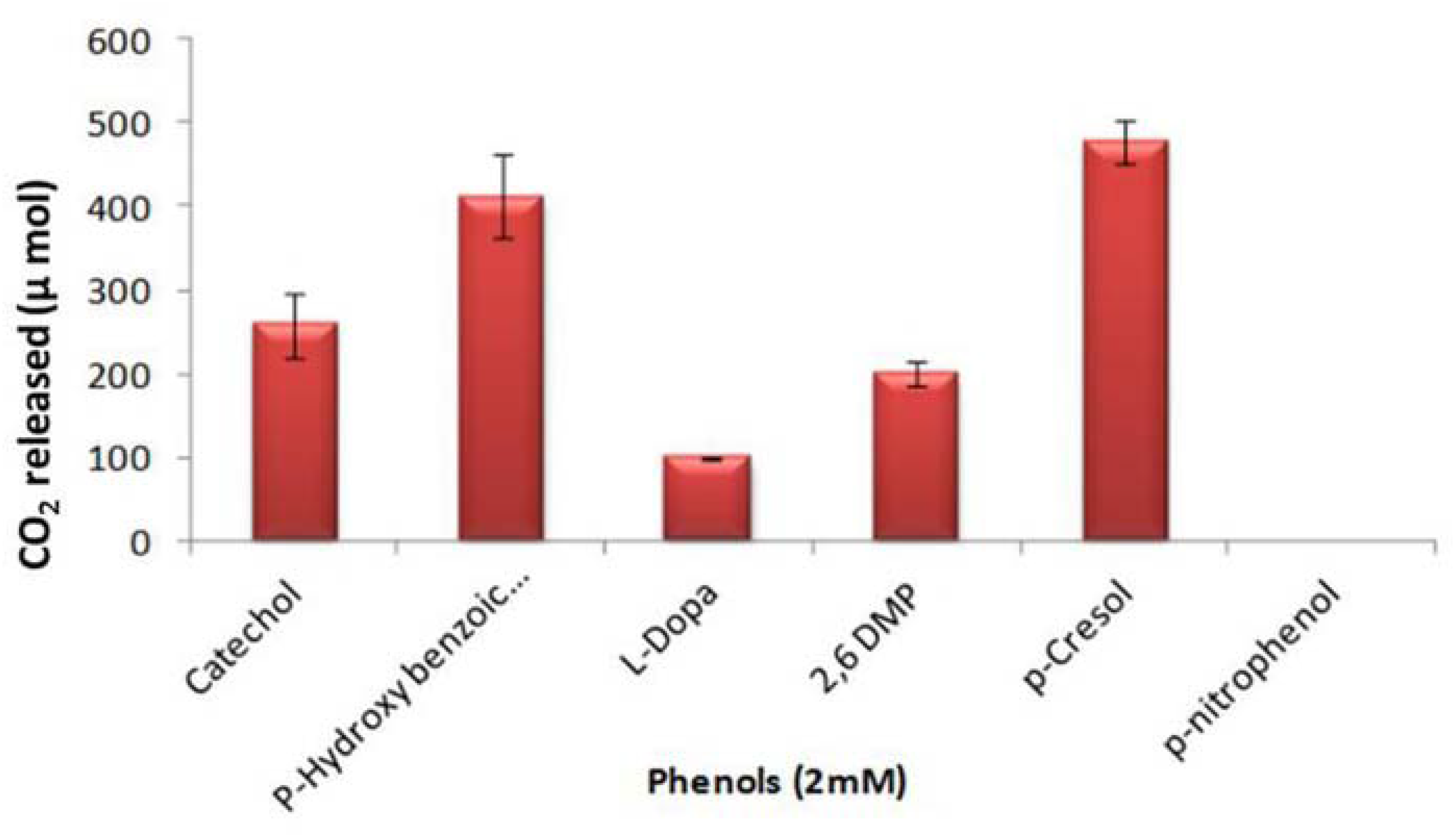
Oxidative polymerization of phenols by TyrB: Mono and di phenols at 2mM concentration with tyrosinase 10 U.ml^−1^ were incubated and the CO_2_ due to oxidative polymerization was monitored. Each phenol was replicated thrice and the values are means ofthree replications. The error bar indicates the standard deviation value.

The results suggest that the enzyme might polymerize the monophenols more efficiently than diphenols that would be released during decomposition of lignin during organic matter decomposition in soil. Although the enzyme could oxidatively polymerize mono and diphenols, *p*-nitro phenol was not oxidized by the enzyme and there was no release of CO_2_.

The reason for this is unclear and needs further study. One of the key reactions in humification is the oxidative polymerization of phenolic compounds which subsequently yield humic substances in the environment [34–36]. Furthermore, the ability of TyrB to generate CO_2_ is directly proportional to cleavage of the ring structure of phenols. Such ring opening of phenols further enables quick degradation which otherwise non-reactive. [35, 37].

The TyrB treatment ends up additional functional groups suggesting the formation of reaction products and confirming the ring cleavage of phenols as observed in the FT-IR spectra. There was no difference in functional groups of p-Nitrophenol and 2,6 Dimethoxy phenol treated and untreated samples. Whereas, the presence of additional two functional groups at wave numbers 1655.59 cm^−1^ corresponding to alkenes C=C medium stretching and 2185.92 cm^−1^ corresponding to alkynes CΞC stretch was noticed in the absorption spectrum of p-cresol. Similarly, the presence of an additional functional group of alkynes CΞC stretch 2147.35 cm-^1^ was noticed for L-DOPA and catechol. Strong aromatic amines at wave number 1327.5 cm-^1^ were noticed for p-Hydroxy benzoic acid treated with TyrB (Table S1). The results of reaction products from ATR-FT-IR show that backbone of humic substances was additionally formed in the tested phenols. It is believed that the alkenes, alkynes and aromatic amines would react with other amino acids present in the environment to form the humic substances[8, 29].

### Transformation of CWW and humification of coir pith biomass

The transformed products obtained from CWW incubation with TyrB (Table S2) revealed that the number of functional groups after incubation was higher in sterilized CWW than the filter sterilized and untreated CWW, indicating the ability of TyrB in oxidative transformation followed by polymerization ability. While comparing the different productsformed during the incubation, it was noticed that the untreated control had only aromatic ring structure which indicates the presence of phenolic group of compounds, whereas similar functional groups with modification were also observed in the sterilized phenols indicating that enzymes were involved in ring cleavage and polymerization[26, 27]. FT-IR spectra of compounds generated from CWW after incubation with TyrB(Fig 4)show the presence of more functional groups in higher intensities in sterilized CWW compared to the control and filter sterilized CWW. Additional functional groups in the wave numbers such as 991.232 cm^−1^, 1078.98 cm^−1^, 1390.42 cm^−1^, and 2085.64 cm^−1^ in higher intensities also indicate that TyrB efficiently transformed CWW(Table S2). On the contrary, filter sterilized CWW didn’t have such additional functional groups (Fig 4).

**Fig 4.**
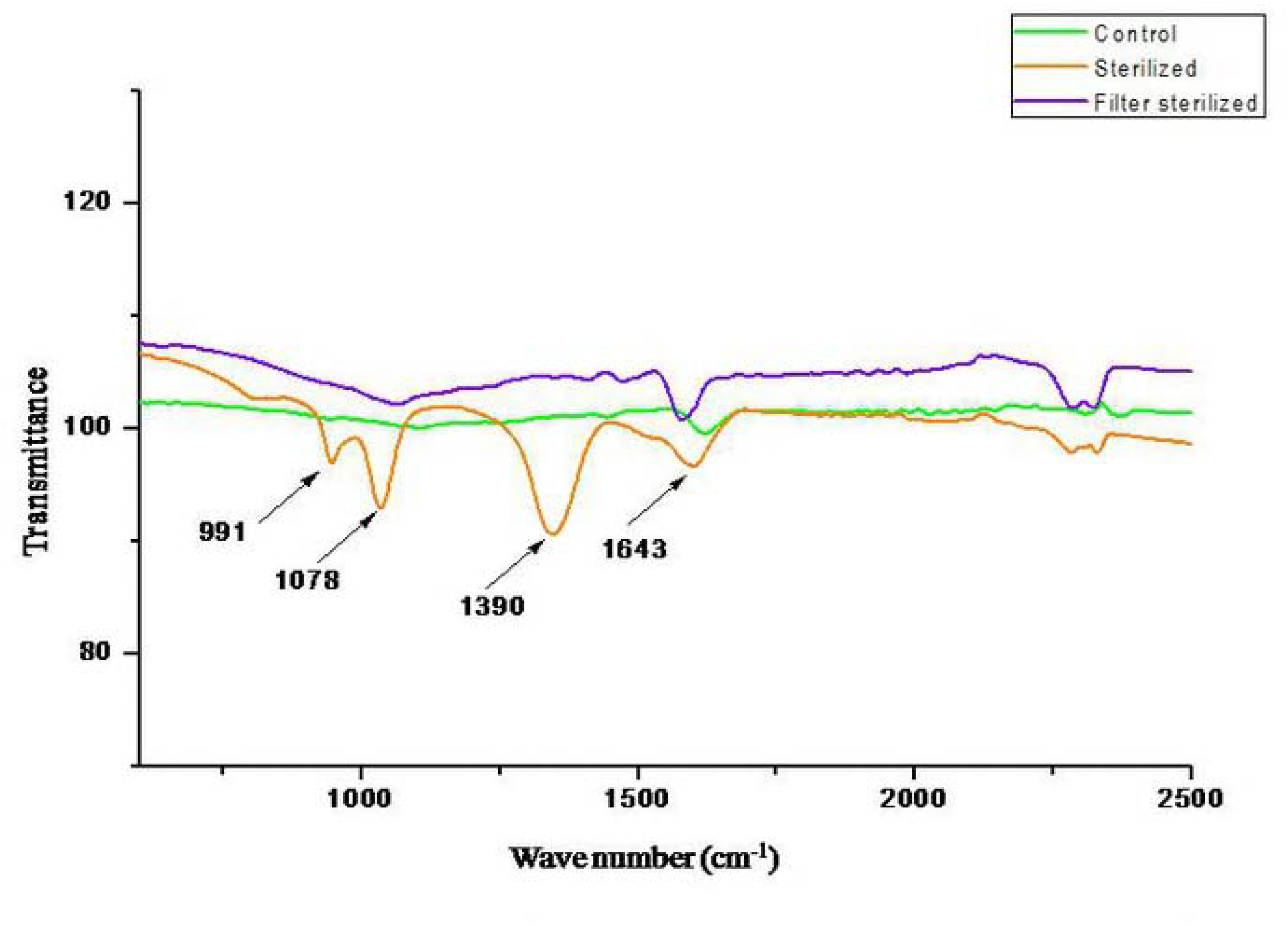
FT-IR spectra of CWW treated with TyrB: FTIR peaks of CWW transformed products of control, sterilized and filter sterilized CWW were presented. Presence of additional peaks at sterilized phenols was indicated by the arrows.

It is believed that in aqueous environment tyrosinase oxidizes low molecular weight phenols and polymerizes them to precipitate it. This enables easy removal of phenolic compounds in wastewater[38]. In the present study, it is evident from the FTIR results that the tyrosinases polymerized phenols as indicated by the presence of additional functional groups.

### Production of humic polymer using coir pith as substrate

Results of UV-Vis spectrum and FT-IR show that the tyrosinase can produce humic substances from coir pith biomass within three days. The UV-Vis spectra of the treated samples exhibited an increase in absorbance over control, and the maximum increment was observed on the 3^rd^ day. The most dramatic increment in absorbance was recorded from 380 to 530 nm (Fig 5). The sudden shift in absorbance at 480 nm was associated with the production of humic substances. Similarly, the increment in absorbance over the three days period was attributed to compounds with carboxylic and phenolic groups as similar pattern of absorbance and molecules were observed when cotton stalk biochar was depolymerized with fungal oxidoreductase[28].

**Fig 5.**
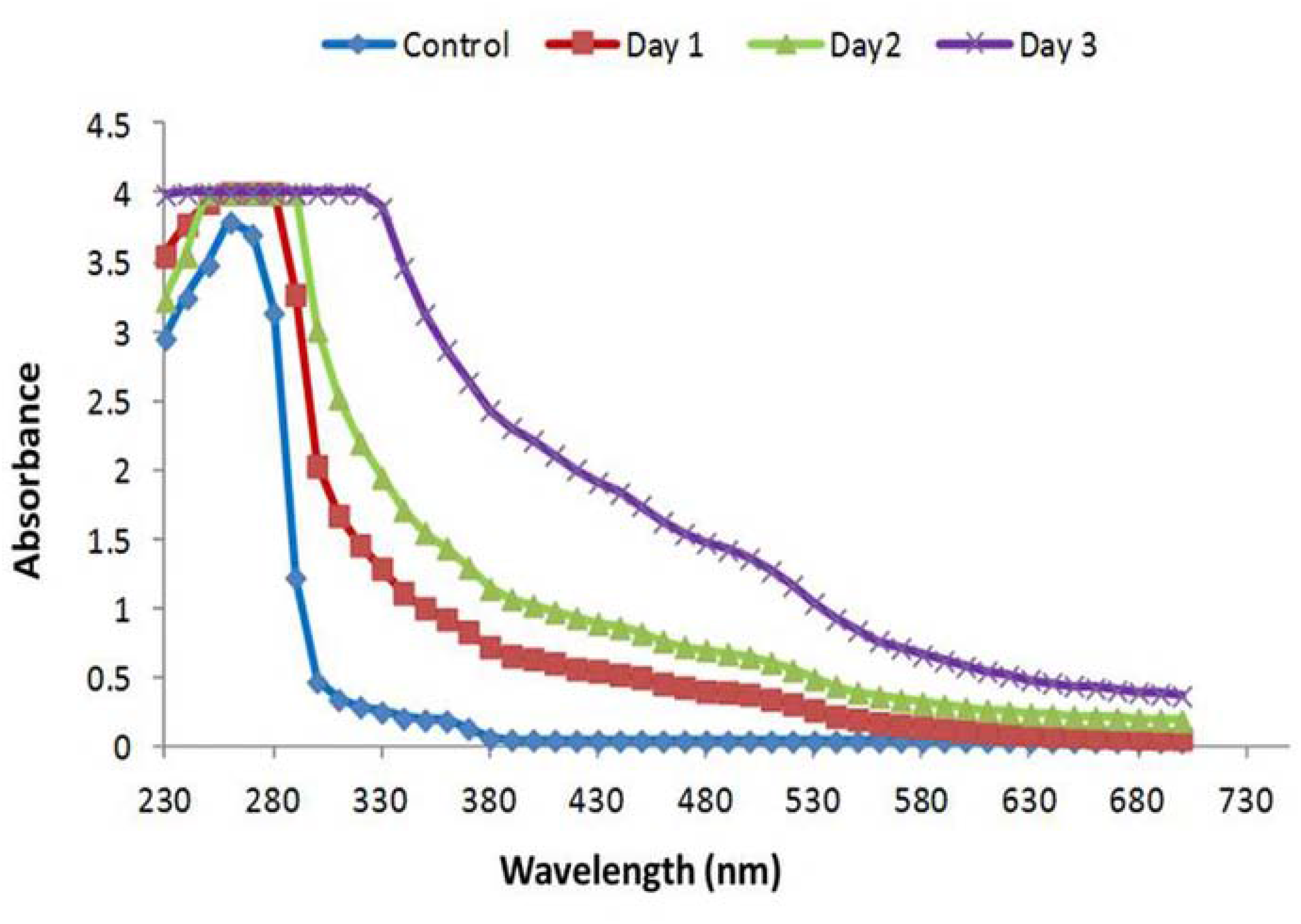
Absorbance spectra or the liquid portion of coirpith biomass after treatment with TyrB: Day wise spectral scan of the treated sterilized CWW showed a hump above 480–530 nm which is related to humic polymer. In addition, several absorbance ratios were calculated for the presence of humic substances and other related products in the liquid portion of the coir pith biomass (Table l).

**Table 1.** UV-Vis absorbance of the liquid phase of coir pith biomass treated with TyrB.

| **Absorbance analysed (nm)** | **Day1** | **Day2** | **Day3** | **Day4** |
| --- | --- | --- | --- | --- |
| 280 | 4.00 | 4.00 | 3.90 | 3.50 |
| 340 | 4.00 | 4.00 | 4.00 | 4.00 |
| 360 | 4.00 | 4.00 | 4.00 | 4.00 |
| 400 | 4.00 | 4.00 | 4.00 | 4.00 |
| 436 | 3.81 | 3.57 | 3.68 | 2.91 |
| 465 | 3.52 | 3.23 | 3.35 | 2.55 |
| 472 | 3.37 | 3.07 | 3.19 | 2.39 |
| 600 | 1.30 | 1.13 | 1.17 | 0.65 |
| 664 | 0.88 | 0.74 | 0.77 | 0.36 |

Maximum absorbance at 280 and 340 nm on all the three days tested is related to the presence of aromatic compounds probably due to depolymerization. Similarly, higher absorbance at 360 until 472 nm indicates the presence of large aromatic compounds like humic substances. Lower values after that do notsignify any compounds related to humic products. Decreasing absorbance over the three days at 436 nm indicates the presence of molecules with more aliphatic, carbohydrates and nitriles compounds (Table 2). The relation between E280/472 describes the presence of aromatic groups in humic substances. The increments in the coefficient on day 2 and the decreased value on day 3 show that depolymerization was obtained by tyrosinase. Similarly, the coefficient E472/664 is associated with the level of condensation of the chain of aromatic carbons. Higher values at this coefficient indicate the presence of more aliphatic structures and less aromatic structures[39–42]. To further study the functional groups associated with humic substances, FT-IR analysis was performed, and the results are presented in Fig6.

**Fig 6.**
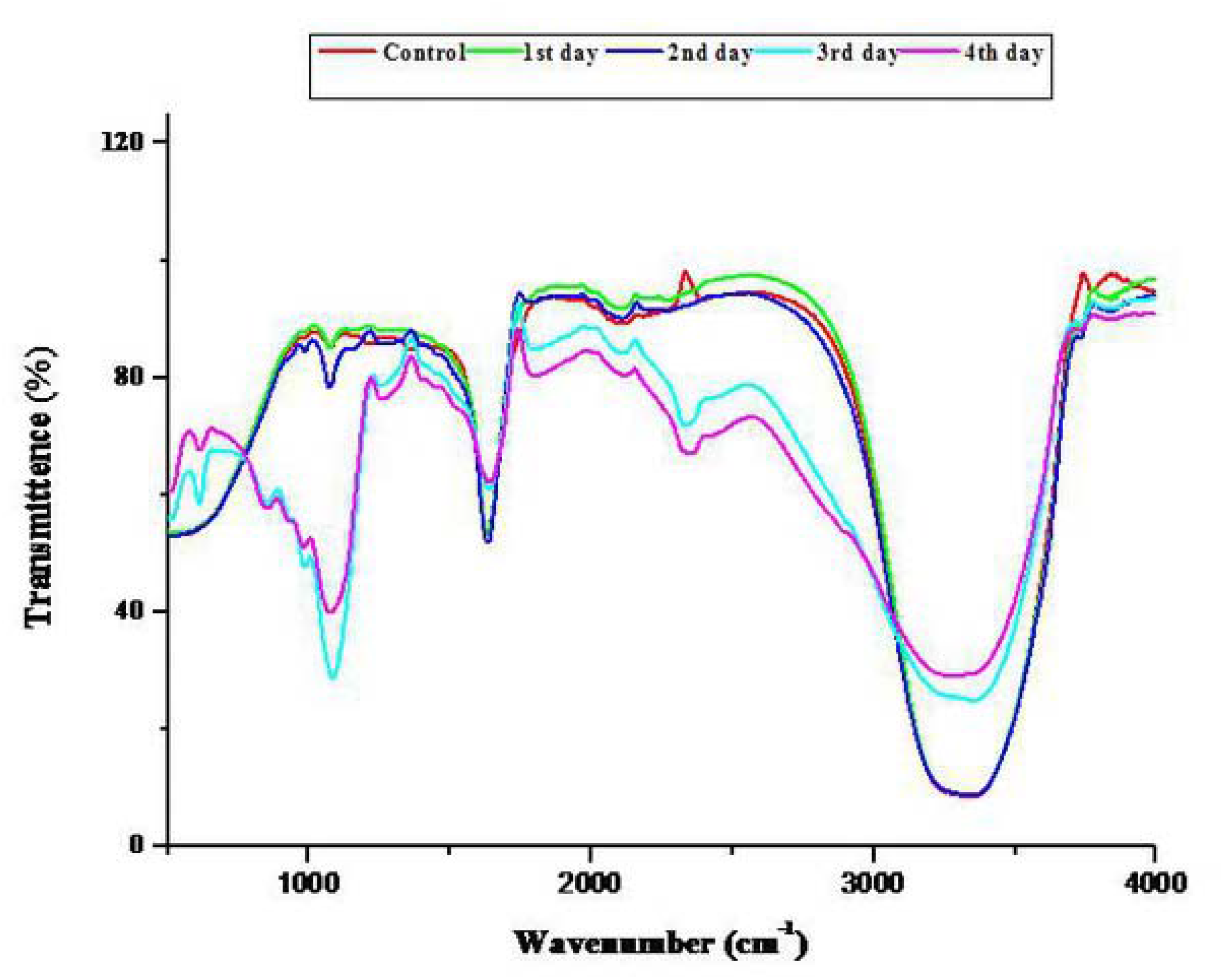
FT-IR spectra of the functional groups of coir pith biomass: Spectral data. were recorded at 64 scans per sec from 400-4000 cm^−1^. Intensities of transmittance: showed increased concentrations of the products released.

**Table 2.** Relationship between absorbance ratios, formation of humic substances and other products in the liquid phase of coir pith biomass.

| **Absorbance Ratios** | **Day1** | **Day2** | **Day3** |
| --- | --- | --- | --- |
| E270/400 | 1.00 | 1.00 | 0.98 |
| E465/665 | 3.99 | 4.34 | 4.31 |
| E250/365 | 1.00 | 1.00 | 0.98 |
| E280/472 | 1.18 | 1.30 | 1.23 |
| E280/664 | 4.51 | 5.36 | 5.02 |
| E472/664 | 3.81 | 4.12 | 4.09 |

The FT-IR spectral data show that aldehyde group(R-CH-O) at wave number 860 cm-^1^ was not observed on the first two days, whereas its intensity was higher on day 3 and reduced on day 4. Similarly, increase in OH groups was recorded as the day progressed and their structural changes as evidenced by changes in vibration were noticed at wave number 1053 cm^−1^. The presence of alkenes at wave number1641 cm^−1^indicates the formation of backbone of humic substances (Fig 6).

There are two possible humification pathway exists in soil as reviewed by [7]. The first theory namely, lignin-protein theory, states that initial material lignin is partially oxidized by oxidases, yielding humic acid which is further oxidized and depolymerized into fluvic acid. In this theory,changes in biomass structure encompasses, loss of OCH_3_ groups from lignin molecule, resulting in the formation of hydroxy phenols, and oxidation of aliphatic chain compounds to form acid groups. Subsequent oxidation of hydroxyl phenols into quinones and semi-quinones, the free radicals formed during the reaction binds to amino and nitrogenous compounds to form HS, This lignin-protein theory are predominant in poorly aerated soils.

The second theory is based on polyphenol theory, which stipulates that during initial lignin breakdown low molecular weight aldehydes and acids are released into the environment. The aldehydes are oxidized into semi-quinones and quinones which undergo non-enzymatic polymerization in the presence of nitrogenous compounds. In this theory, fluvic acids are formed prior to humic acid. Polyphenol theory is more common in well-aerated forest soils.

In comparison of two theories with the present study, FT-IR analysis of coir pith biomass after TyrB treatment shows that aldehydes and acids were present initially in higher concentrations on the third day (Fig 6). This signifies that breakdown products of lignin by TyrB are present. Further their absence on fourth day onwards suggests that the aldehydes and acidic groups might have undergone polymerization as evidenced by an increase in absorbance on the 3^rd^ day [Fig 5 and Table 1]. Therefore, based on the spectroscopic studies (UV-VIS, FT-IR),the structural changes and release of compounds during humification of coir pith biomass suggest that TyrB might follow polyphenol theory of HS formation as reported by [7].

## Conclusion

The enhanced catalytic bio-transformation and formation of HS from coir pith biomass by TyrB produced by *B.aryabhattai* TFG5 clearly suggest that the TyrB follows polyphenol theory of humification and has a potential for application in phenolic industry. In addition, the oxidative transformation of CWW into products would enable this enzyme to be used in detoxification and precipitation of toxic phenols in coir industry. The HS produced through this process could be a slow-release organic fertilizer in organic agriculture which can enhance the carbon sequestration potential of soil.

## Acknowledgements

Financial support granted to SU from Department of Biotechnology, Government of India (No.BT/PR4891/BCE/8/905/2012 & 17.07.2013) is gratefully acknowledged.

## Supporting Information

**S1 Fig.** Multiple sequence alignment of TyrB. MSA was performed using ClustalW program using Bioedit v. 7.2.5. The tyrosinase sequences from the various organisms and their accession numbers are given inside ear bracket are *Bacillus aryabhattai* (WP_043981293.1), *B. megaterium* (3NM8), *Bacillus* sp. Root147 (WP_057234811), *B. flexus* (WP_025750930), Fictibacillusmacauensis (WP_050979754), B. macauensis ZFHKF-1 (EIT84795),*Pseudomonas veronii(WP_017849537), Nitrosomonas europaea* ATCC 19718 (CAD85152), *P. fluorescens* (WP_047297073), *Streptomyces tsukubensis* (WP_040914590), *Streptomyces* sp. NRRL S-87 (WP_030192477), *Brevibacillussp*. is *Brevibacilluslaterosporus* GI-9 (CCF17084), *S. roseus* (WP_048477298),*S. roseoverticillatus*(WP_030366439). The copper binding His sites and Phe site are boxed. The color shades represent identical and conserved amino acids in all the organisms.

**S1 Table.** FTIR reaction products from the phenols and the formation of additional functional 370 groups.

**S2 Table.** FT-IR functional groups and their corresponding wavenumber of CWW.

